# Nasal delivery of single-domain antibodies improves symptoms of SARS-CoV-2 infection in an animal model

**DOI:** 10.1101/2021.04.09.439147

**Authors:** Kei Haga, Reiko Takai-Todaka, Yuta Matsumura, Tomomi Takano, Takuto Tojo, Atsushi Nagami, Yuki Ishida, Hidekazu Masaki, Masayuki Tsuchiya, Toshiki Ebisudani, Shinya Sugimoto, Toshiro Sato, Hiroyuki Yasuda, Koichi Fukunaga, Akihito Sawada, Naoto Nemoto, Chihong Song, Kazuyoshi Murata, Takuya Morimoto, Kazuhiko Katayama

## Abstract

The severe acute respiratory syndrome coronavirus 2 (SARS-CoV-2) that causes the disease COVID-19 can lead to serious symptoms, such as severe pneumonia, in the elderly and those with underlying medical conditions. While vaccines are now available, they do not work for everyone and therapeutic drugs are still needed particularly for treating life-threatening conditions. Here, we showed nasal delivery of a new, unmodified camelid single-domain antibody (VHH), termed K-874A, effectively inhibited SARS-CoV-2 titers in infected lungs of Syrian hamsters without causing weight loss and cytokine induction. *In vitro* studies demonstrated that K-874A neutralized SARS-CoV-2 in both VeroE6/TMPRSS2 and human lung-derived alveolar organoid cells. Unlike other drug candidates, K-874A blocks viral membrane fusion rather than viral attachment. Cryo-electron microscopy revealed K-874A bound between the receptor binding domain and N-terminal domain of the virus S protein. Further, infected cells treated with K-874A produced fewer virus progeny that were less infective. We propose that direct administration of K-874A to the lung via a nebulizer could be a new treatment for preventing the reinfection of amplified virus in COVID-19 patients.

**Author summary:** Vaccines for COVID-19 are now available but therapeutic drugs are still needed to treat life-threatening cases and those who cannot be vaccinated. We discovered a new heavy-chain single-domain antibody that can effectively neutralize the severe acute respiratory syndrome coronavirus 2 (SARS-CoV-2) that causes COVID-19. Unlike other drug candidates, which prevent the virus from attaching to the receptor on the host cell, this new antibody acts by blocking the virus membrane from fusing with the host cell membrane. We studied the behavior of the new antibody *in vitro* using VeroE6/TMPRSS2 cells and human lung organoids. When delivered through the nose to infected Syrian hamsters, we found that this antibody could prevent the typical symptoms caused by SARS-CoV-2. Our results are significant because delivering simple drugs directly to infected lungs using a nebulizer could increase the potency of the drugs while reducing the risk of immune reaction that could occur if the drugs escape or are delivered through the blood stream.

## Introduction

Coronaviruses (CoV) are enveloped, single-stranded positive RNA viruses. They are divided into four genera: alpha, beta, gamma and delta. Betacoronaviruses are significant interest because they are responsible for the severe acute respiratory syndrome (SARS) and middle eastern respiratory syndrome (MERS) epidemics in the past and now, the coronavirus disease 2019 (COVID-19) pandemic. These viruses can cause mild to severe respiratory tract infections and death in some cases. All betacoronaviruses feature a spike protein on their surface that assists entry into cells. The spike protein has two distinct subunits, S1 and S2, and the receptor binding domain (RBD) in the S1 subunit interacts with the host cell receptor. Because the identity of the spike protein in SARS-CoV-1 (virus responsible for SARS) and SARS-CoV-2 (virus causing COVID-19) is 76% similar, both viruses bind to the same host cell receptor, angiotensin-converting enzyme 2 (ACE2). After binding to ACE2 via the RBD domain, a protease on the host cell surface cleaves and activates the S protein, allowing the virus membrane to fuse with the host cell membrane. Blocking this viral fusion is thought to be a promising therapeutic strategy [1, 2]. Furthermore, because SARS-CoV-2 infects cells in the nasal mucosa or lungs that express ACE2 [3], direct delivery of antiviral drugs to the respiratory system is expected to improve drug efficacy.

Camelid single-domain antibodies are a unique class of antibodies that consist of single heavy chain. Variable domain of heavy chain of heavy chain antibodies (VHHs) have been developed to bind viruses such as influenza virus, human immunodeficiency virus 1 (HIV-1) and respiratory syncytial virus (RSV) [4]. These antibodies, which consist of single amino acid chains, can bind specific antigens with high affinity and specificity. Unlike human monoclonal antibodies, VHHs can be easily modified and produced using bacteria. They are also stable against heat and pH, allowing them to be stored longer than human monoclonal antibodies and therefore, stockpiled for epidemics. Being stable also means VHHs can be nebulized and administered via an inhaler directly to infected lungs. The first VHH undergoing clinical trials for direct delivery into the lung by nebulization to treat RSV is ALX-0171 [4]. *In vitro* studies show ALX-0171 binds to the F protein in RSV with higher affinity than approved human monoclonal antibody prophylactic drug, Palivisumab [5]. Intranasal administration to cotton rats reduced RSV load in the nose and lung [5, 6]. More recently, VHH against SARS-CoV-1 and SARS-CoV-2 have also been identified [7, 8]. This VHH (termed VHH072), which must fuse with an Fc domain of a human antibody to work against SARS-CoV-2 [7], neutralizes the virus by binding to the RBD in the S protein. While effective *in vitro*, exactly how VHH072 neutralizes SARS-CoV-2 and how it performs *in vivo* remain unclear.

Here, using the S1 domain of SARS-CoV-2 S protein as an antigen, we screened an extensive DNA library and found a standalone VHH that is specific for SARS-CoV-2. This new VHH (termed K-874A) bound SARS-CoV-2 S protein with higher affinity than previous VHHs [7, 8] and does not require any modification with antibody fragments, making them a very attractive therapeutic candidate. We show K-874A could neutralize SARS-CoV-2 effectively in VeroE6/TMPRSS2 and human normal alveolar-derived cells. Different from other therapeutics, which act by blocking viral attachment to ACE2, K-874A neutralizes SARS-CoV-2 by preventing viral membrane from fusing with the host cell membrane. Cryo-electron microscopy analysis revealed that K-874A binds to both the RBD and N-terminal domain (NTD) regions of the S protein rather than at the interface of the RBD and ACE2 host receptor. Studies in human lung-derived alveolar organoid showed that infected cells treated with K-874A produced fewer virus progeny that were also less infective, suggesting that K-874A can block the virus from spreading to uninfected cells and persons. When intranasally administered to SARS-CoV-2-infected Syrian hamsters, K-874A could prevent weight loss, reduce viral replication in the lungs and prevent cytokine storms that are characteristic of a severe SARS-CoV-2 infection. Our results demonstrate the K-874A is a potent therapeutic drug against SARS-CoV-2.

## Results

### *In vitro* selection of anti-SARS-CoV-2 S1 VHHs

We used VHH-cDNA display for *in vitro* selection of VHHs against the SARS-CoV-2 S1 protein (Fig 1A). cDNA display yields functional VHHs, whose coding RNA is linked via a puromycin linker [9-11]. This procedure selects the coding sequences for high-affinity VHHs from a diverse (10^13-14^) DNA library [12]. To select the VHH candidates targeting SARS-CoV-2 S1 proteins, DNA libraries at Round 2 (R2) and Round 3 (R3) of *in vitro* selection were sequenced, and anti-SARS-CoV-2 VHH candidates were translated.

**Fig 1.**
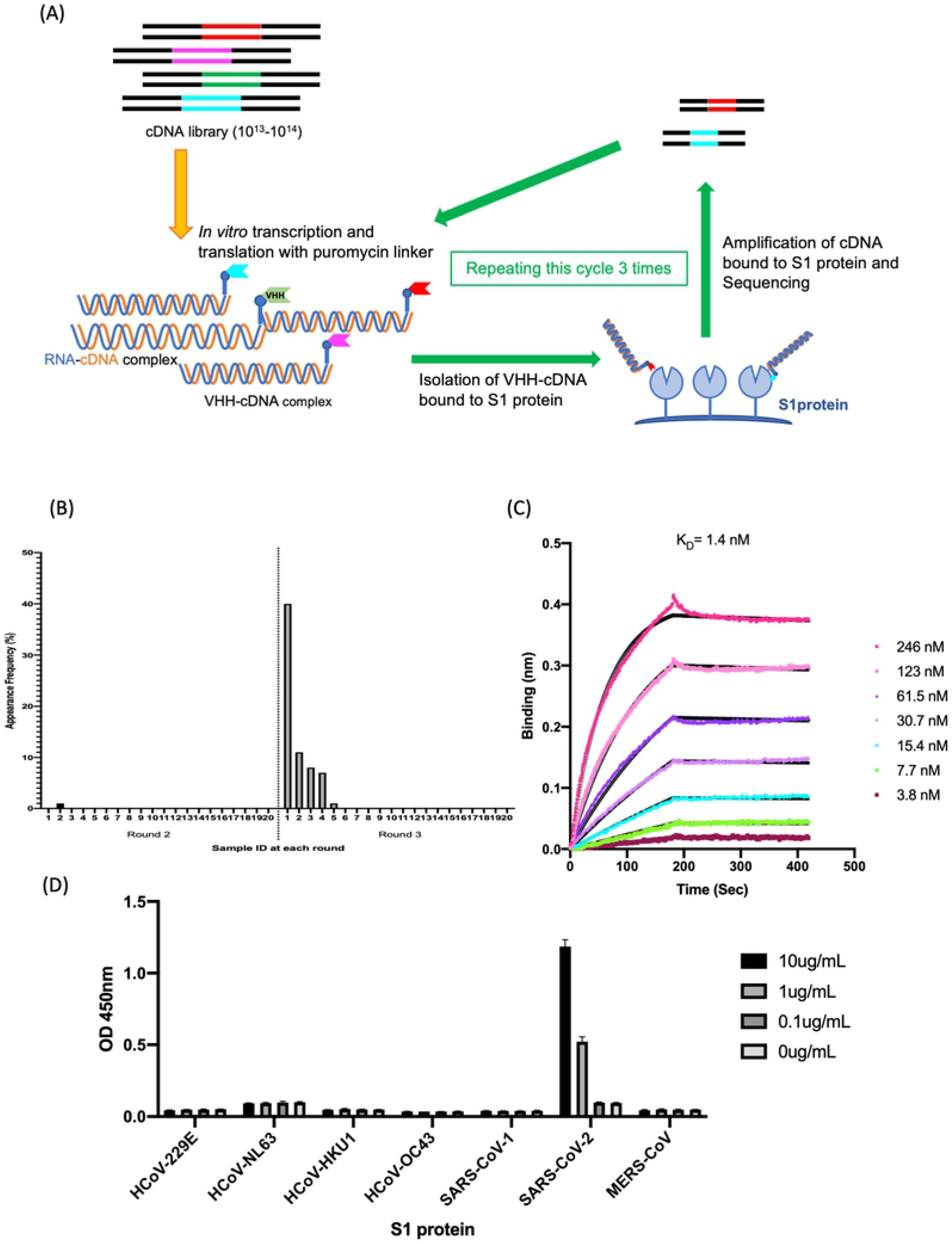
Isolation and characterization of K-874A. (**A**) Schematic showing *in vitro* selection of VHHs against SARS-CoV-2 S1 protein using VHH-cDNA display. *In vitro* transcription and translation of VHH-cDNA library forms VHH linked to its mRNA with puromycin linker. cDNA of linked mRNA was revers transcribed and VHH-cDNA complex was produced. High-affinity VHH-cDNA complex to immobilized S1 protein was isolated and its cDNA was amplified. Three rounds of selection were performed and cDNA libraries from rounds 2 and 3 were sequenced and anti-SARS-CoV-2 VHH candidates were translated. (**B**) Frequency distribution of amino acid sequences corresponding to VHH antibody candidates targeting SARS-CoV-2 S1 subunits in the selected VHH libraries. Sample ID “1” with the highest frequency (39.5%) is clone K-874A. (**C**) Binding affinity of K-874A to SARS-CoV-2 S1 subunits. Biolayer interferometry sensorgram measures the apparent binding affinity of K-874A-6xHis to immobilized SARS-CoV-2 S1 fused with sheep Fc. Binding curves for different concentrations of K-874A are shown in different colors. Black curve is 1:1 fit of the data. (**D**) Direct antigen ELISA measuring the apparent binding affinity of FLAG-tagged K-874A to immobilized S1-6xHis subunits of alpha- (HCoV-229E and HCoV-NL63) and beta-coronaviruses (HCoV-HKU1, HCoV-OC43, SARS-CoV-1, SARS-CoV-2 and MERS CoV). Error bars are mean ± S.D. (N=3). Data are from an experiment representative of 3 independent experiments.

From the frequency distribution of VHH clones appearing in the selected library, clone K-874A appeared most frequently (39.5%) after three rounds of *in vitro* selection, indicating that K-874A has a high affinity to S1 proteins (Fig 1B). Biolayer interferometry assay showed the binding affinity of K-874A to SARS-CoV-2 S1 protein is 1.4 nM (K_a_ (1/Ms) = 6.72E+04, K_d_ (1/s) = 9.42E-05) (Fig 1C). When a direct antigen enzyme-linked immunosorbent assay (ELISA) was performed using FLAG-tagged K-874A and immobilized recombinant His-tagged S1 protein, we found that K-874A bound with high affinity to the S1 protein of SARS-CoV-2 but not to other coronaviruses including human CoV (HCoV-229E, HCoV-NL63, HCoV-HKU1, HCoV-OC43), MERS-CoV and SARS-CoV-1 (Fig 1D). Together, these results indicate that K-874A binds strongly and specifically to the S1 protein of SARS-CoV-2.

### K-874A VHH prevents SARS-CoV-2 fusion with host cell

With its strong binding affinity and specificity to SARS-CoV-2, we investigated K-874A as a potential therapeutic drug for COVID-19. We infected African green monkey kidney (VeroE6) cells expressing transmembrane protease serine 2 (VeroE6/TRMPSS2) with SARS-CoV-2 and measured how well K-874A inhibited SARS-CoV-2 infection in the cells using quantitative real-time polymerase chain reaction (qRT-PCR). Half-maximal inhibitory concentration (IC_50_) calculated from RNA copies was 1.40 μg/mL (Fig 2A). IC_50_ indicates the concentration of K-874A required to inhibit 50% of SARS-CoV-2.

**Fig 2.**
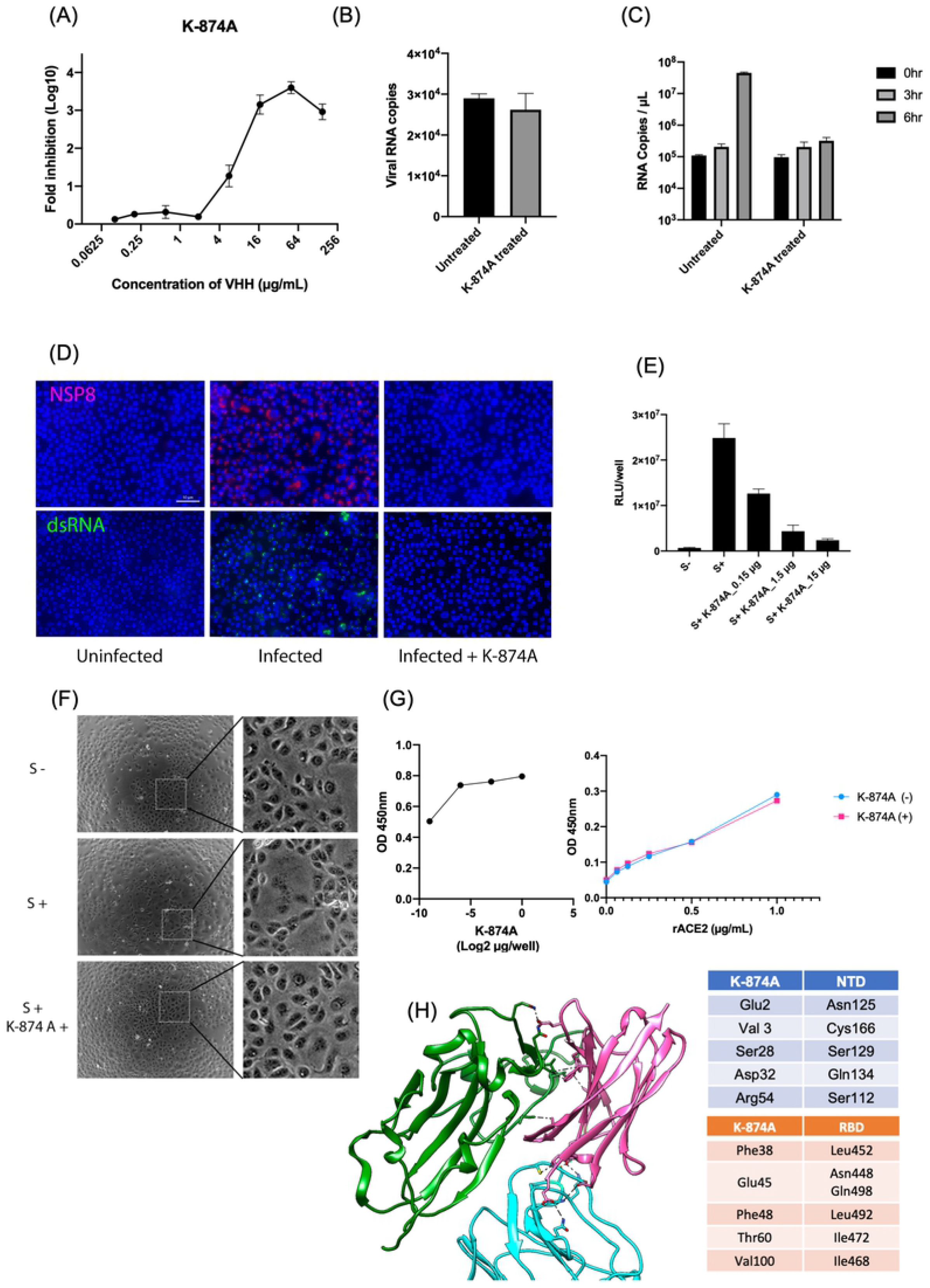
K-874A VHH neutralizes SARS-CoV-2 by preventing viral fusion with host cell. (**A**) Neutralization of SARS-CoV-2 by K-874A VHH. SARS-CoV-2 was pretreated with different concentrations of K-874A and inoculated into VeroE6/TRMPSS2 cells. Fold-reductions in viral RNA copies in culture supernatants were calculated by qRT-PCR. K-874A has an IC_50_ of 1.40 μg/ml. Error bars indicate mean ± SD from three independent experiments. (**B**) Viral RNA copies in VeroE6/TMRPSS2 cells infected with SARS-CoV-2 (untreated) and SARS-CoV-2 pretreated with K-874A VHH (K-874A-treated) were similar. Number of RNA copies attached or incorporated in the cells were determined by qRT-PCR. Error bars indicate mean ± SD from three independent experiments. (**C**) Number of viral RNA copies in VeroE6/TMRPSS2 cells infected with untreated SARS-CoV-2 and K-874A-treated SARS-CoV-2 at 0, 3 and 6 hours. RNA was extracted from the cells and estimated by qRT-PCR. (**D**) Immunofluorescence images show viral protein, NSP8, and RNA replication are suppressed in Vero/TMPRSS2 cells infected with K-874A-treated SARS-CoV-2. SARS-CoV-2-infected cells were fixed at 24 hours after infection and treated with anti-NSP8 (red) or anti-dsRNA (green). After fluorescence-conjugated secondary antibody and Hoechest (blue) treatment, images were capture by BZX800. Scale bar = 50 μm. (**E**) Cell fusion induced by S protein transduction in HiBiT- and LgBiT-expressing VeroE6/TMPRSS2 cells. Treatment with K-874A VHH suppressed cell fusion in a dose-dependent manner. Error bars indicate mean ± SD from three independent experiments. (**F**) Optical images of VeroE6/TMPRSS2 cells expressing S protein transduced by a lentivirus vector show cell fusion (S+) and no cell fusion when treated with K-874A VHH (S+K-874A+). Inset: magnified view of dotted square shown in left panels. (**G**) ELISA results show K-874A (left panel) and ACE2 (right panel) bound to immobilized recombinant S protein in a dose-dependent way. Immobilized S protein was detected by serially diluted K-874A (1 to 0.002 μg/well) and horse radish peroxidase (HRP)-conjugated anti-VHH antibody (left panel), or by serially diluted ACE2 protein (1 to 0.0625 μg/ml) and anti-ACE2 antibody and HRP-conjugated secondary antibody (right panel). Binding of ACE2 to S protein was unaffected by K-874A (1μg/well) treatment. (**H**) Homology docking model of S protein and K-874A. Amino acid residues were predicted by auto refinement using Phoenix. NTD and RBD of S protein are shown in blue and green, respectively. K-874A is in magenta. Interacting residues are in black dotted lines and summarized in the right panel.

To determine whether K-874A VHH neutralizes SARS-CoV-2 by preventing the virus from attaching to the host cell, we compared the early phases of infection in cells infected with K-874A-treated virus and untreated virus. VeroE6/TMPRSS2 cells were incubated with K-874A-treated SARS-CoV-2 and untreated virus for 1 hr and the number of RNA copies that are attached or incorporated into the cells were determined by qRT-PCR. K-874A treatment did not change the levels of virus attachment or early incorporation in the cells (Fig 2B). At 6 hours post-infection, the number of viral RNA copies in cells infected with K-874A-treated virus remained unchanged while the numbers were significantly higher in cells infected with untreated virus (Fig 2C). Additionally, expression of the viral protein, NSP8, and double stranded RNA (dsRNA), which is generated during viral RNA replication, were suppressed in cells infected with K-874A-treated viruses (Fig 2D). These results indicated that K-874A did not reduce viral attachment.

We further investigated whether K-874A acts by inhibiting the virus from fusing with the host cell. VeroE6/TMPRSS2 cell expresses the ACE2 receptor and the TMPRSS2 protease on its surface. To study viral fusion, we transduced the VeroE6/TMPRSS2 cells with a lentivirus coding for the viral S protein into the cells. When the S protein from one VeroE6/TMPRSS2 cell binds to the ACE2 receptor on an adjacent VeroE6/TMPRSS2 cell, the TMPRSS2 protease cleaves and activates the S protein for fusion. To quantify this cell fusion, we used HiBiT technology, which involves the binding of HiBiT (a small 11–amino acid peptide) with a larger subunit, called LgBiT, to form a complex with luciferase activity. VeroE6/TMPRSS2 cells expressing HiBiT or LgBiT were produced and co-cultivated. The HiBiT-LgBiT complex and the resulting luciferase activity form only when the S protein from one cell binds to the ACE2 receptor on an adjacent cell and their membranes are fused. We found that treating VeroE6/TMPRSS2 cells with K-874A suppressed cell fusion in a dose-dependent way (Fig 2E). Cells without S protein (S -) did not fuse while those expressing S protein but were not treated with K-874A (S+) did fuse. Optical micrographs confirmed these results (Fig 2F). Direct binding of K-874A and recombinant ACE2 to immobilized recombinant S protein in ELISA further showed that both K-874A and ACE2 bound to S protein in a dose-dependent way (Fig 2G). However, K-874A did not block ACE2-S protein interaction (right panel in Fig 2G). Together, these findings suggest that K-874A does not prevent the virus from attaching to the ACE2 receptor on the cell surface. Rather, K-874A prevents the virus from entering the cell by blocking the viral membrane from fusing with the host cell.

To estimate the K-874A-binding region on the S protein, we compared the cryo-electron microscopy (cryo-EM) structures of recombinant S trimer and the K-874A-S trimer complex. In the prefusion state, the RBD is known to move upwards to bind ACE2 [13, 14]. We observed that K-874A was located in the vacant space between the NTD and RBD of the S protein in the prefusion conformation (Fig 2H). CDR1 and CDR2 amino acids and the N-terminal of K-874A formed polar bonds with the NTD of the S protein while CDR2 and CDR3 bonded hydrophobically with the RBD. These findings demonstrate that K-874A neutralizes SARS-CoV-2 via a different route that did not involve ACE2 binding.

### K-874A VHH prevents viral replication in lung organoids

Because SARS-CoV-2 targets lung tissues and induces severe respiratory disease, we evaluated K-874A VHH in a human lung-derived alveolar organoid, which we have shown is susceptible to SARS-CoV-2 and releases high levels of progeny virus into the culture supernatant[15] (Ebisudani T *et al*. submitted). To evaluate whether K-874A can block the production of virus progeny in infected alveolar organoid, we incubated the organoid cells with SARS-CoV-2 for 1 hr before treating the infected cells with K-874A-containing medium for 3 days. Untreated cells released >1×10^5^ RNA copies/μL into the culture supernatant at day 3, whereas VHH treatment reduced the progeny production to 1×10^4^ RNA copies/μL (Fig 3A), indicating that K-874A can reduce the production of virus progeny. To test the infectivity of virus progeny produced from K-874A-treated and untreated infected cells, we incubated the virus progeny with fresh VeroE6/TMPRSS2 cells. Untreated virus progeny showed 78.4 TCID_50_/μL, while K-874A-treated virus progeny showed less than 1.2 TCID_50_/μl (Fig 3B). These data suggested that the K-874A-treated progeny have a lower infectivity. This is likely due to K-874A VHH binding to the S protein of the virus progeny, preventing them from infecting other cells.

**Fig 3.**
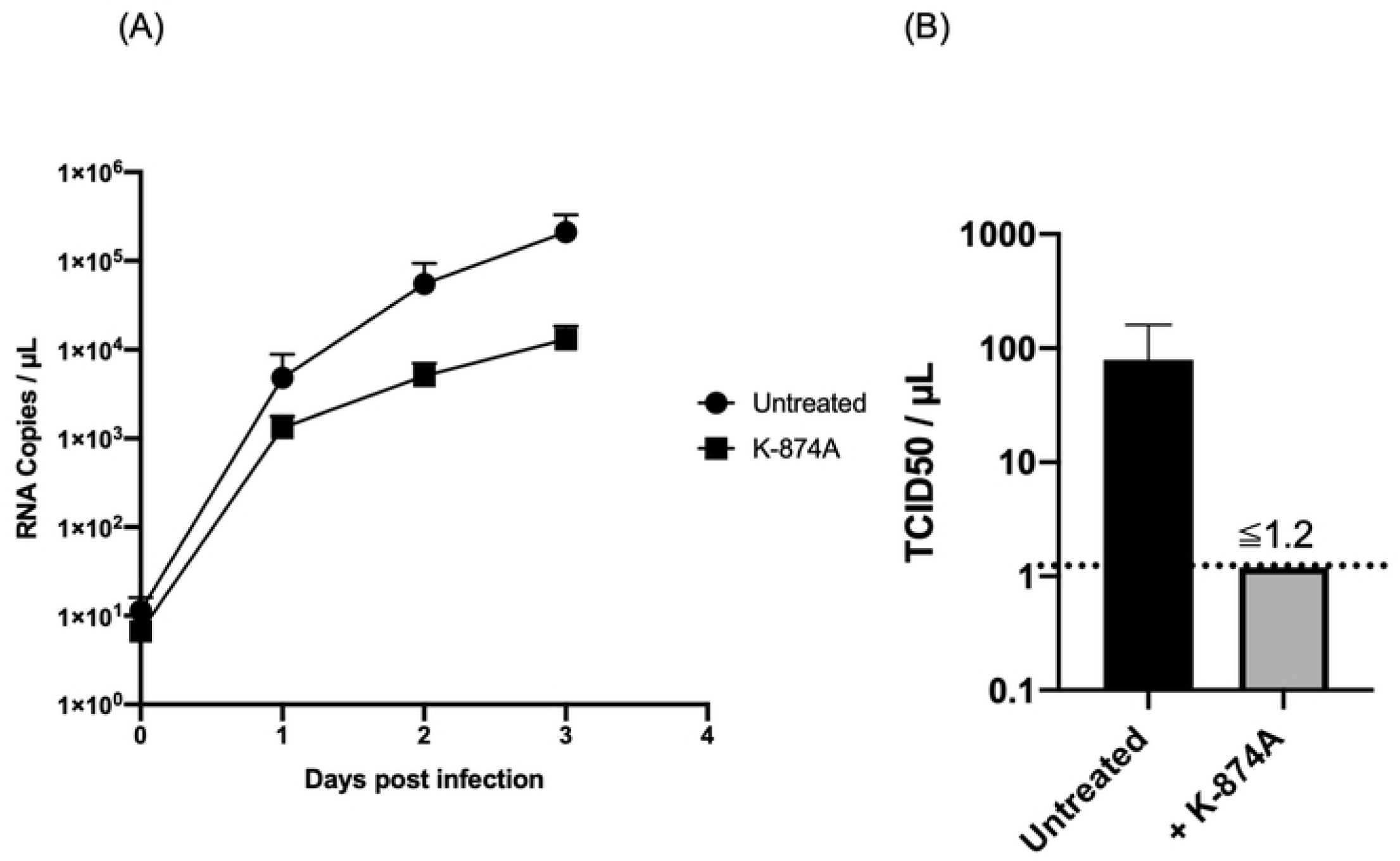
K-874A VHH treatment inhibits SARS-CoV-2 replication in an alveolar organoid model. (**A**) Graph shows infected alveolar organoid treated with K-874A-containing medium for 3 days released fewer RNA copies (1×10^4^ RNA copies/μL) into the culture supernatant than organoids that were not treated with K-874A (>1×10^5^ RNA copies/μL). (* P<0.005, One-way ANOVA using a Dunnett’s test). Data are from an experiment representative of 2 independent experiments. (**B**) Virus progeny in culture supernatant of K-874A-treated infected alveolar organoid had a lower infectivity (<1.2 TCID_50_/μl) than progeny from untreated organoids (78.4 TCID_50_/μL) at Day 2. Virus infectivity was determined in VeroE6/TMPRSS2 cells. Error bars indicate mean ± SD. Black dotted line indicates detection limit.

### Infected Syrian hamsters recover after K-874AVHH treatment

We further tested K-874A VHH as a potential therapeutic drug for COVID-19 using the Syrian hamster animal model. When infected with SARS-CoV-2, Syrian hamsters lose weight but spontaneously heal over time [16]. To determine whether K-874A can improve the symptoms of COVID-19 infection, we infected 6 week-old male Syrian hamsters with SARS-CoV-2 (2 x 10^3^ TCID_50_) and intranasally administrated different regimens of K-874A. The animals were treated with either a single dose of K-874A just before the infection or at 1 day after the infection, or multiple doses at 1 and 2 days after the infection (Fig 4A). Each dose was 3 mg K-874A per animal. All infected animals treated with K-874A did not lose weight whereas infected control animals treated with bovine serum albumin in phosphate buffered saline (BSA/PBS) exhibited weight loss 3 days after the infection (Fig 4B). Further, all K874A-treated hamsters regardless of treatment regimen had a statistically lower number of viral RNA copies in their lung tissues than BSA/PBS-treated hamsters 4 days after the infection (Fig 4C). Because cytokines are known to be upregulated after an infection [17, 18], we compared the mRNA levels of several cytokines in the infected lung tissues of the untreated and K-874A-treated animals. While untreated animals showed elevated levels of IL6 and IL10 but not IL1β and TNF-α, in their lungs 4 days after the infection, all K-874A-treated animals regardless of the treatment regimen displayed cytokine levels that were similar to uninfected animals (Fig 4D).

**Fig 4.**
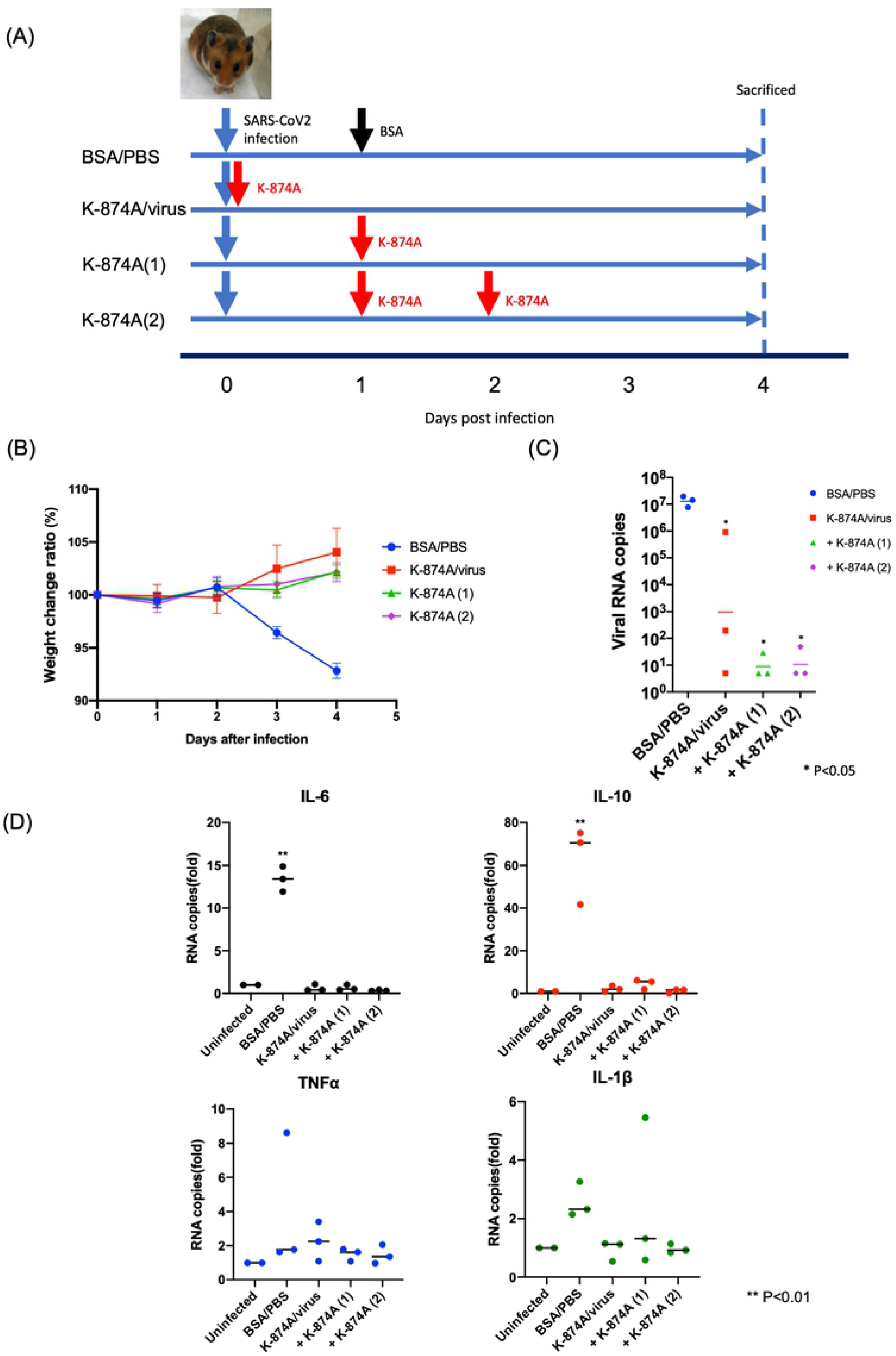
K-874A VHH treatment improves viral symptoms in Syrian hamsters. (**A**) Time schedule for inoculation and K-874A administration to Syrian hamsters. All animals were infected with 2×10^3^ TCID_50_ of SARS-CoV-2. Untreated animals were treated with BSA/PBS (Blue circle) on Day 1 and Day 2 post-infection. K-874A-treated groups were treated with either a single dose immediately before infection (K-874A/virus; Red square) or 1 day after infection (+ K-874A (1); Green triangle), or two doses at Day 1 and Day 2 after infection (+ K-874A (2); Purple rhombus). Each dose was 3 mg K-874A per animal. (N=3 for infected with untreated or treated) (**B**) Unlike untreated animals, all K-874A-treated animals showed no weight loss. Weight at Day 0 is 100%. (**C**) Amounts of viral RNA in the lung homogenates at Day 4 as determined by qRT-PCR show all K-874A-treated animals had a statistically lower number of viral RNA copies in their lungs than untreated animals. (**D**) qRT-PCR measurements of lung homogenates collected on Day 4 show untreated animals expressed elevated levels of inflammatory cytokine IL-6 and IL-10 but not TNFα and IL-1β. Cytokine levels in K-874A-treated animals were similar to uninfected animals. (N=2 for uninfected)

When different doses (0.12 mg, 0.6 mg or 3 mg per animal) of K-874A were administered in a two-dose regimen (on Day 1 and Day 2 post-infection), animals receiving the lowest K-874A dose displayed the highest weight loss and viral RNA copies in their lung tissues. Their IL-6 and IL-10, but not IL1β and TNF-α, levels were also elevated compared to those receiving higher doses of K-874A (Supplementary Fig 2). These data demonstrate that K-874A can improve the symptoms of COVID-19 infection and it acts in a dose-dependent way.

## Discussion

Besides vaccines, therapeutic drugs are much needed for the treatment of COVID-19 infections. VHH are promising drug candidates because they are more stable and cheaper to produce than human monoclonal antibodies. They are also amenable to nasal administration, allowing high concentrations of drugs to reach directly to infected lungs and remain effective for longer. Indeed, nasal administered VHH against RSV have been shown to be effective for 3 days [6]. However, due to their low molecular weight, VHH monomers have very short half-lives in the blood stream [4]. VHHs under development as antivirals need to be either made multivalent or modified with human antibody fragments to enhance their antiviral effect or extend their half-life. A previous report showed VHH that bound to the S protein of SARS-CoV-2 could neutralize the virus only when VHH was fused with an Fc domain of a human antibody [7].

Here, using *in vitro* selection, we identified a standalone anti-SARS-CoV-2 S1 VHH that binds to the S protein of SARS-CoV-2 with a higher affinity than previous VHHs [7, 8] and displays excellent neutralizing ability in VeroE6/TMPRSS2 cells and human normal alveolar-derived cells. Our results show that this VHH neutralizes SARS-CoV-2 by preventing the virus membrane from fusing with the host cell membrane. Cryo-EM analysis of the S protein-VHH complex revealed that the VHH binds between the RBD and NTD region on the S protein, rather than at the interface of the RBD and ACE2. Studies in human lung-derived alveolar organoid indicated that the VHH can reduce the production of virus progeny. Moreover, virus progeny produced from VHH-treated infected cells had a lower infectivity than those that came from untreated infected cells, suggesting that the VHH could prevent the virus from spreading to uninfected cells or persons. Intranasal administration of our VHH to SARS-CoV-2-infected Syrian hamsters prevented weight loss, viral replication in the lungs and the upregulation of cytokines typically caused by SARS-CoV-2 infection.

Our VHH has several advantages. A previous study showed that inhaling a human monoclonal antibody against SARS-CoV-2 can inhibit virus replication in the lung and nasal turbinate [19]. This suggests that the therapeutic benefits of our VHH could also be delivered using a nebulizer. Nasal administration via a nebulizer is expected to lower the amount of VHH entering the blood stream, which could reduce the risk of immunoreaction against VHH on repeated use. Further, because our VHH displays striking antiviral effects without the additional Fc domain, its risk to Fc-related antibody-dependent enhancement is likely to be low.

Abnormal secretion of cytokines is a hallmark of serious cases of COVID-19 infections [20]. While it remains unknown why such cytokine storms occur, it has been suggested that ACE2 may be an interferon-stimulated gene that SARS-CoV-2 could exploit to enhance infection [3]. Because our VHH prevents viral fusion, we believe that it can reduce viral infection and in turn, inhibit interferon upregulation. The efficacy of our VHH could also be improved if used in conjunction with antibodies that block ACE2-RBD binding.

## Acknowledgements

This work was supported by the Japan Agency for Medical Research and Development (AMED No. 20fk0108295h0001), by Platform Project for Supporting Drug Discovery and Life Science Research (Basis for Supporting Innovative Drug Discovery and Life Science Research (BINDS)) from AMED (No. 2833), and the COVID-19 Kitasato project. We thank Akikazu Murakami for providing the VHH library and Ai Lin Chun of Science Storylab for critically reading and editing the manuscript.

## Author contributions

KH, TM, and KK conceived the project. KK acquired funding. KH, TTR, TTakano, CS, KM, and AS conducted the investigation. KH, TTR, TTakano, YM, TM, KK, NN, MT, HM, TToujou, AN, and YI contributed methods. TM and KK administered the project. TTakano, YM, TM, TE, SS, HY, KF, TS, KK, and TToujou provided resources. KH, TTR, and YM provided visualization. KH, TTR, and YM wrote the original draft. KH, TM, TS, and KK reviewed and edited the paper.

---

**Supplementary figure 1.**
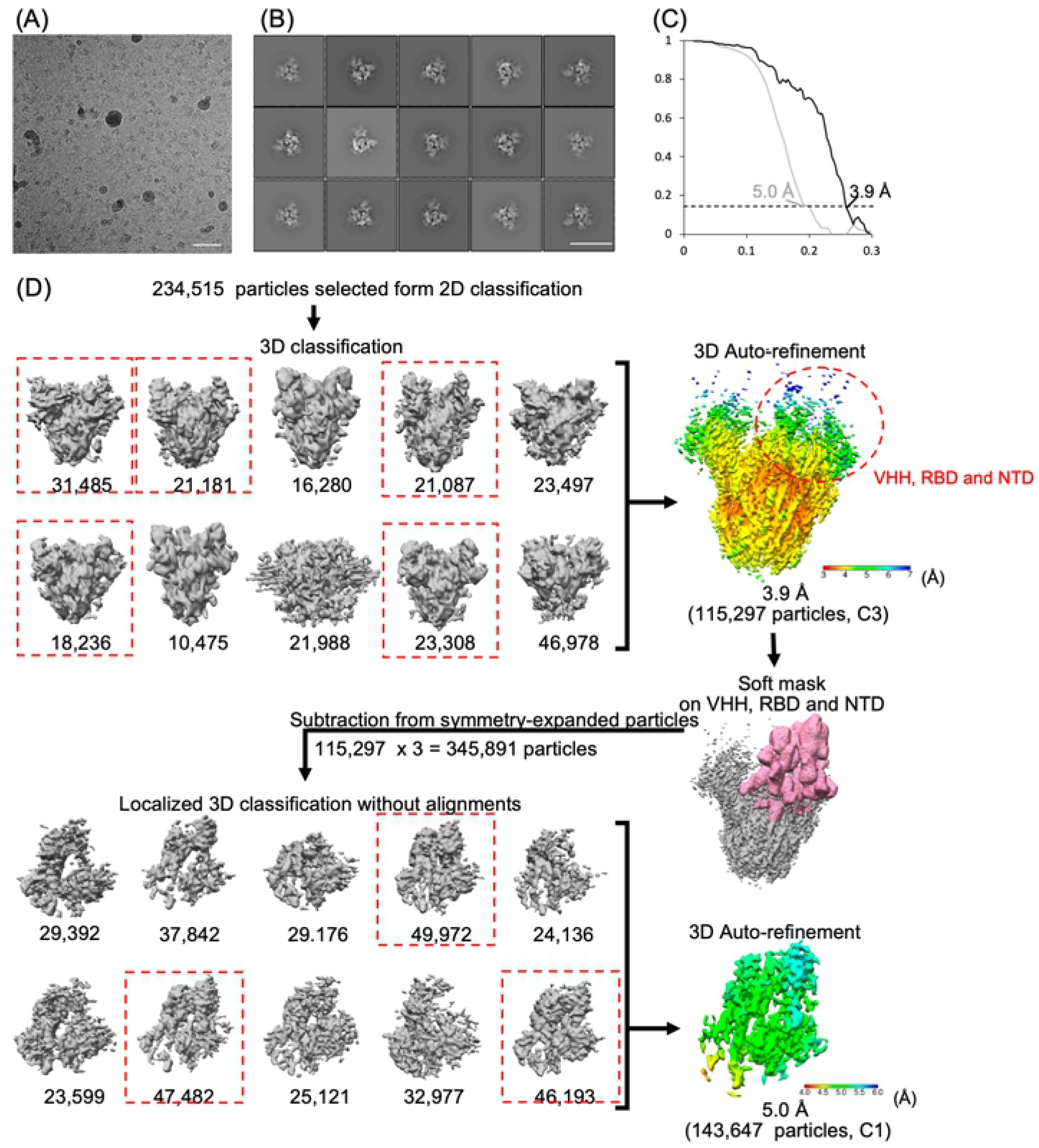
Cryo-EM analysis of S protein trimer of SARS-CoV-2 with VHH antibodies. (A) A representative motion-corrected electron micrograph. Scale bar, 5 nm. (B) Representative 2D class averaged images. Scale bar, 2nm (C) Gold-standard Fourier shell correlation (FSC) curves for the 3D maps of the overall structure (black) and the structure focused on VHH, RBD and NTD. black curve (grey). (D) The workflow for cryo-EM data processing.

**Supplementary figure 2.**
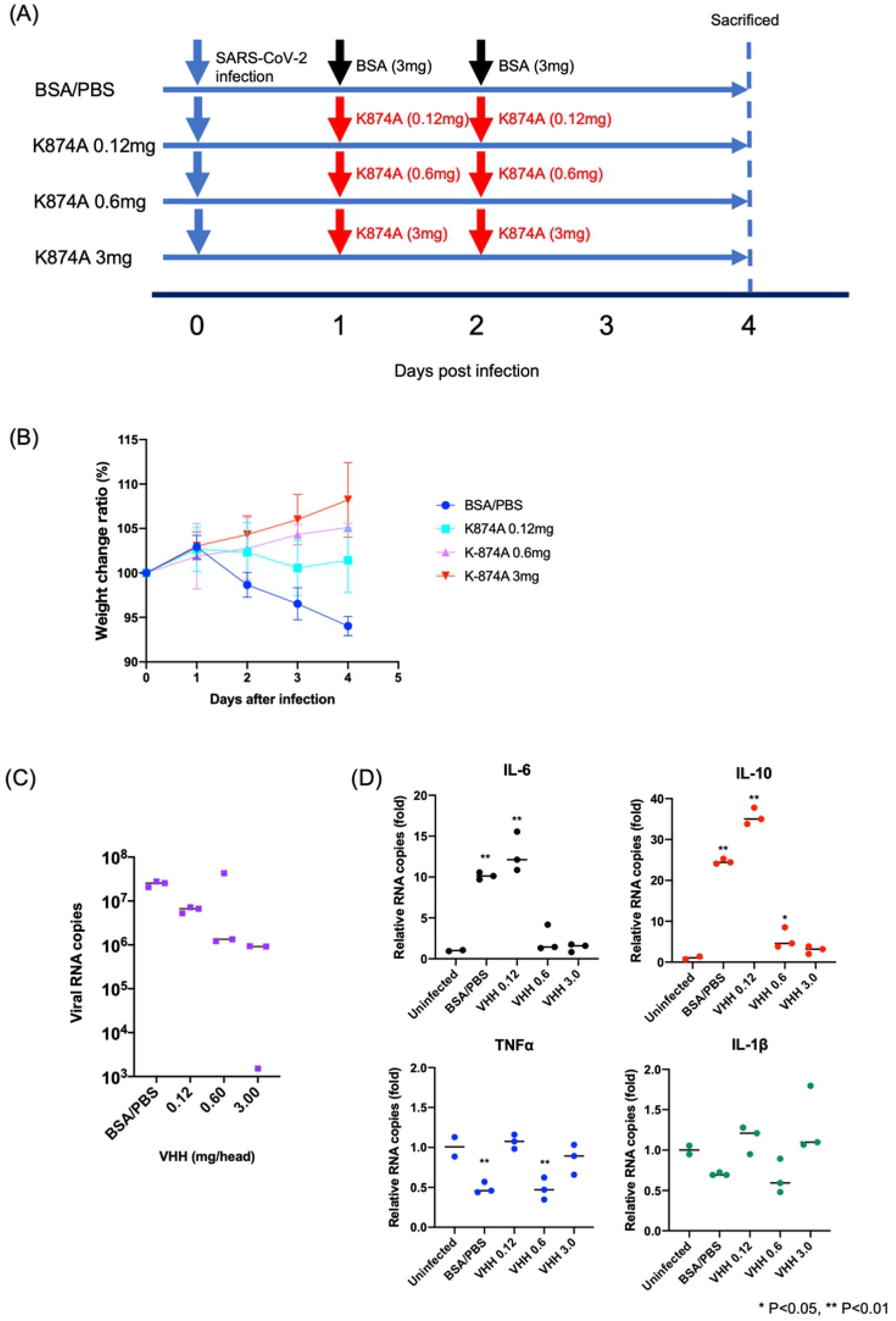
Different dose of VHH Treatment to infected Syrian Hamsters. (**a**) Time schedule for inoculation and K-874A administration to Syrian hamsters. **(b)** Weight changes in K-874A-treated and untreated hamsters after SARS-CoV-2 infection as indicated. Weight at Day 0 is 100%. **(c)** Amounts of viral RNA in the lung homogenates at Day 4 were determined by qRT-PCR. (**d**) qRT-PCR results showing inflammatory cytokine expression. Expression levels of IL-6, IL-10, TNFα and IL-1β were assessed in lung homogenates collected at Day 4.

## EXPERIMENTAL PROCEDURES

### Preparation of cDNA display library

A cDNA display library (200 pmol-scale) used for the 1^st^ round of selection (R1) was synthesized from an initial VHH-encoding DNA library (predicted diversity, 1×10^13^), provided by Epsilon Molecular Engineering Co. Ltd. (Japan). A cDNA display library (6 pmol-scale) used at the 2^nd^ and 3^rd^ rounds of selection (R2 and R3, respectively) was synthesized from the selected DNA library of each previous round. Translation of the DNA library was performed using T7 RiboMAX™ Express Large-Scale RNA Production System (Promega, USA), according to the manufacturer’s instructions. The DNase-treated mRNA mixture was purified using Agencourt RNAClean XP beads (Beckman Coulter Genomics, USA). The purified mRNA was hybridized to cnvK (riboG) linker (Epsilon Molecular Engineering) at the 3’ -terminal region in 50 mM Tris-HCl (pH 7.5) with 200 mM NaCl under the following annealing conditions: heating at 90°C for 2 min, followed by lowering the temperature to 70°C at a rate of 0.1°C/s, incubating for 1 min, then cooling to 25°C at a rate of 0.1°C/s, incubating for 30 sec., and then stored at 4°C until use. The mixture was irradiated with UV light at 365 nm using a handheld UV lamp (6W, UVGL-58, 254/365 nm, 100V; Analytik Jena, USA) for 5 min to obtain mRNA-linker complex.

The mRNA-linker complex was translated using a Rabbit Reticulocyte Lysate System, Nuclease-Treated (Promega) at 37°C for 15 min. To synthesized mRNA-linker-VHH complex, KCl and MgCl_2_ were added to final concentrations of 900 and 75 mM, respectively, and the mixture was incubated at 37°C for 20 min.

EDTA (final concentration: 70 mM) and an equal volume of 20 mM Tris-HCl buffer (pH 8) containing 2 mM EDTA, 2 M NaCl and 0.2% Tween 20 were added, and the mRNA-linker-VHH complex was immobilized in 60 μL of Dynabeads MyOne streptavidin C1 magnetic beads (ThermoFisher Scientific, USA) at 25°C for 30 min. The beads were washed two times with 60 μL of wash buffer composed of 10 mM Tris-HCl (pH 8), 1 mM EDTA, 1 M NaCl and 0.1% Tween 20. The immobilized library was then reverse-transcribed by ReverTra Ace (Toyobo Life Science) at 42°C for 30 min. The beads were washed two times with 200 μL of wash buffer composed of 20 mM sodium phosphate buffer, 0.5 M NaCl, 5 mM imidazole, and 0.05% Tween 20 (pH 7.4), and then 30 μL of RNase T1 (ThermoFisher Scientific) prepared with the wash buffer was added to the beads. The mixture was incubated for 15 min 37°C to release the cDNA-linker-VHH complex from the beads. The supernatant containing cDNA-linker-VHH complex was purified using His Mag Sepharose Ni beads (GE Healthcare, USA). The supernatant was added to 20 μL of the beads and incubated at 25°C for 30 min. The beads were washed with 200 μL of wash buffer composed of 20 mM sodium phosphate buffer, 0.5 M NaCl and 20 mM imidazole (pH 7.4), and then incubated in 10 μL of elution buffer composed of 20 mM sodium phosphate buffer, 0.5 M NaCl and 250 mM imidazole, 0.05 % Tween 20 (pH 7.4) at 25 °C for 15 min. The elution collected was stored at 16°C until use.

### *In vitro* selection of VHHs using cDNA display

SARS-CoV-2 S1 subunits-His tag (Sino Biological, China) were used as target molecules for screening. Microtiter wells of Nunc-Immuno™ Plate II (ThermoFisher Scientific) were coated overnight at 4°C with 100 μL of 100 μg/mL (R1), 10 μg/mL (R2), 1 μg/mL (R3) recombinant SARS-CoV-2 S1 subunits-His tag which was prepared with PBS. The S1 subunits-coated wells were blocked with 200 μL of 3% BSA in PBST for 2 hours at room temperature, and then washed at three times with 200 μL of HBST. 100 μL of VHH-cDNA complex library was prepared as follows: 50 μL of HEPES buffer containing 1% BSA was added to 50 μL of VHH-cDNA conjugate library for R1, 50 μL of HBST containing 1% BSA, 25 μL of HBT was added to 50 μL of cDNA display library for R2 and R3, respectively. The VHH-cDNA complex library was added to the target-fixed well, incubated for 1 hour at room temperature. The residual VHH-cDNA conjugates were removed by washing at 10 times with 200 μL of HBST. The target-binding VHH-cDNA complex was eluted by adding 100 μL of 100 mM Tris (hydroxymethyl) aminomethane (pH 11), gently pipetting, and incubating for 10 min at 37°C. The elution was immediately transferred to PCR cocktails prepared with KAPA HiFi HotStart ReadyMix (2x) (Kapa Biosystems, USA), forward primer (5 ‘-GATCCCGCGAAATTAATACGACTCACTATAGGGGAAGTATTTTTACAACAATTACCA ACA-3’), and reverse primer (5’-TTTCCACGCCGCCCCCCGTCCT-3’). The PCR cycle conditions consisted of 95°C for 2 min, followed by 22 cycles (R1) or 24 cycles (R2 and R3) of 98°C for 20 sec, 68°C for 15 sec, and 72°C for 20 sec, and then 72°C for 5 min. The PCR products were purified using Agencourt AMPure XP beads (Beckman Coulter Genomics), according to the manufacturer’s instruction. The purified DNA libraries were used for translation, as described above, to prepare cDNA display libraries used at the next selection round.

### Sequencing of the DNA library

DNA libraries obtained at R2 and R3 were used as templates in amplicon PCR for Illumina sequencing. The DNA library was quantified using PicoGreen^®^ dsDNA reagent kit (ThermoFisher Scientific) and prepared at 100 ng/mL with nuclease-free water. 1^st^ PCR was performed with 12.5 μL of KAPA HiFi HotStart ReadyMix (2x) (Kapa Biosystems), 0.5 μL of 10 μM forward primer (5 ‘-TCGTCGGCAGCGTCAGATGTGTATAAGAGACAGNNNNATGGCTGAGGTGCAGCTCG TG-3 ‘), 0.5 μL of 10 μM reverse primer (5 ‘-GTCTCGTGGGCTCGGAGATGTGTATAAGAGACAGNNNNTGATGATGATGGCTACCACCTCCCG-3’), 9 μL of nuclease-free water, and 2.5 μL of template DNA. The PCR cycle conditions consisted of 95°C for 3 min, followed by 16 cycles of 98°C for 20 sec, 62°C for 15 sec, and 72°C for 20 sec, and then 72°C for 5 min. PCR clean-up was performed with Agencourt AMPure XP beads (Beckman Coulter Genomics). Subsequently, index PCR was performed with 12.5 μL of KAPA HiFi HotStart ReadyMix (2x) (Kapa Biosystems), each 1 μL of 10 μM forward and reverse primers from Nextera XT Index Kit v2 (Illumina, USA), 8 μL of nuclease-free water, and 2.5 μL of template DNA. The PCR cycle conditions consisted of 95°C for 3 min, followed by 8 cycles of 98°C for 20 sec, 55°C for 15 sec, and 72°C for 30 sec, and then 72°C for 5 min. After magnetic bead-based purification of PCR products, the library concentration was measured with PicoGreen^®^ dsDNA reagent kit (ThermoFisher Scientific) and prepared to 4 nM with nuclease-free water. The library was diluted to a final concentration of 7 pM and 5% of PhiX DNA (Illumina, USA) was added. Sequencing was performed with the Miseq Reagent Nano kit V2 500 cycle using a MiSeq 2000 (Illumina), according to the manufacturer’s instructions.

### Data analysis

Dell Mobile Precision 7520 (Xeon E3-1535M v6 (Quad core, 3.1 GHz, 4.2 GHz turbo), 1TB SSD, memory: 64 GB (4×16 GB), 2,400 MHz DDR4 ECC SDRAM) was used to analyze sequencing data. The raw Illumina paired-end reads that passed through the Q30 filter were merged using PEAR software [21]. with default parameter on Linux. Nucleotide sequences encoding VHHs were extracted using combination of basic Linux command lines specifying typical nucleotide sequences conserved on VHHs frame regions. The VHHs-encoding sequences were translated based on standard genetic code using MEGA X software [22].

### Purification of VHHs

*Bacillus subtilis* 168, which is deficient for nine proteases (i.e., *epr, wprA, mpr, nprB, bpr, nprE, vpr, aprE* and *aprX*)[23] and contains a sigma factor for a sporulation (*sigF*)-deficient mutant was used (JP4336082B2). *B. subtilis* was precultured in L medium (1% tryptone, 0.5% yeast extract, and 0.5% NaCl) and cultured in 2xL-Mal medium (2% tryptone, 1% yeast extract, 1% NaCl, 7.5% maltose hydrate, and 7.5 μg/mL MnSO_4_) with 15 μg/mL tetracycline at 30°C 72 h.

Supernatants were subjected to SDS-PAGE and western blotting. SDS-PAGE used SuperSep™ Ace, 15-20% Tricine Gel (Wako, Japan), and then proteins in the gel were transferred to PVDF membrane using Trans-Blot Turbo Mini PVDF Transfer Packs (Bio-Rad, USA) and Trans-Blot Turbo System (Bio-Rad). PVDF membranes were treated with 6xHis Tag Monoclonal Antibody (3D5) HRP (Invitorogen, USA) for 6xHis-tagged proteins, and ANTI-FLAG M2-peroxidase (HRP)-conjugated (Sigma Aldrich, USA) for FLAG^®^ (DYKDDDDK)-tagged proteins in iBind Western System (Invitrogen). Each tagged protein was visualized by 1-Step Ultra TMB-Blotting Solution (ThermoFisher Scientific).

CoVHH1-6xHis and CoVHH1-CoVHH1-6xHis were purified from the supernatants of culture medium using Ni-NTA agarose beads (Wako) and resuspended into PBS with 50 mM Imidazol. The CoVHH1-FLAG fraction was collected from the supernatants using an Amicon^®^ Ultra Centrifugal Filter Unit (Merck, USA) and resuspended into PBS with 50 mM imidazol. The purified VHHs were stored at 4°C until use.

### ELISA for evaluation of the binding specificity of CoVHH1

Microtiter wells of Nunc-Immuno™ Plate II (ThermoFisher Scientific) were coated overnight at 4°C with 100 μL of recombinant human-CoV-229E, NL63, HKU1, and OC43, and MARS-CoV, SARS-CoV-2 S1 subunits-His tag (Sino Biological) and recombinant SARS-CoV S1 subunits-His tag (The Native Antigen Company, UK) diluted in phosphate-buffered saline (PBS) containing 0.05% Tween 20 (PBST). The S1 subunits-coated wells were blocked with 200 μL of 5% skim milk in PBST for 1 hour at room temperature, and then washed at three times with 200 μL of PBST. 100 μL of 20 μg/mL CoVHH1-FLAG prepared with PBST were added to the wells, incubated for 1 hour, and then washed at three times with 200 μL of PBST. The binding of CoVHH1-FLAG to S1 subunits-His tag was detected by incubating for 1 hour at room temperature with mouse monoclonal ANTI-FLAG^®^ M2-Peroxidase (Merck) diluted with PBST. The wells were washed three times with PBST, and then 100 μL of substrate buffer prepared with an o-phenylenediamine dihydrochloride substrate tablet (ThermoFisher Scientific), 10x Stable Peroxide Substrate Buffer (ThermoFisher Scientific), and water. After incubating for 30 min at room temperature, the absorbance at 450 nm was immediately measured with Microplate Reader Infinite M1000 PRO (TECAN, Switzerland).

### Biolayer interferometry

Binding affinities of CoVHH1 and S1 protein were determined using biolayer interferometry (BLI) of Octet RED 384 (Pall Life Sciences, USA). The kinetic buffer was PBST for BLI. The temperature was fixed at 25°C. 50 μL of each measurement solution was added to a 384-well black plate (Fortebio, USA), and the measurements were performed as below. 1) The loading baseline was measured in kinetic buffer for 30 sec. 2) CoVHH1-6xHis was immobilized on a HIS1K biosensor (Fortebio) and loaded until a 0.3-nm signal was achieved. 3) The measurement baseline was measured in kinetic buffer for 30 sec. 3) The loaded sensors were dipped into twofold serial dilutions from 245.8 nM of the SARS-CoV-2 (2019-nCoV) Spike S1-Fc Recombinant Protein (Sino Biological) to measure a 180-sec specific binding at the association step. 4) The dissociation was obtained by dipping the biosensors once more time into the kinetic buffer for 240 sec. The data were analyzed using ForteBio Octet analysis software (Date Analysis HT Version 11.1.2.48) (Fortebio) and kinetic parameters were determined using a 1:1 monovalent binding model.

### Cells and virus

VeroE6/TMPRSS2 were purchased from JCRB Cell Bank and preserved in our laboratory. SARS-CoV-2, 2019-nCoV JPN/TY/WK-521 were provided from National Institute of Infectious Diseases (NIID, Japan)

### Neutralization assay

2.5×10^3^ TCID_50_ of SARS-CoV-2 (MOI=0.05) was incubated with serial diluted VHHs for 2 hours at 37°C and subsequently for 24 hours at 4°C. VHH-treated virus was incubated with 5.0×10^4^ of VeroE6/TMPRSS2 cells for 1 hour at 37°C, and the wells were washed to remove unbound viruses. At 24 hours after infection, the culture supernatant was used to determine the number of RNA copies of virus by quantitative real-time PCR (SARS-CoV-2 Detection Kit, TOYOBO), according to manufacturer’s protocol. IC_50_ was determined 3 days after infection.

### SARS-CoV-2 binding on and early replication in Vero/TMPRSS2 cells

2.5×10^4^ TCID_50_ (MOI=50) of SARS-CoV-2 was incubated with 150 μg/ml of VHHs for 2 hours at 37°C and then 24 hours at 4°C and inoculated to 5.0×10^4^ VeroE6/TMPRSS2 cells for an hour. Similarly, to compare early RNA replication in the infected cells, infected cells were collected at 3 or 6 hours after infection. Total RNA was extracted from the collected cells with nucleospin (Takara). Numbers of RNA copies of the virus were determined by quantitative real-time PCR (SARS-CoV-2 Detection Kit, TOYOBO), according to manufacturer’s protocol.

### Cell fusion assay

To quantify the cell fusion induced by S protein, HiBiT technology (Promega) was applied. HiBiT was linked to the C-terminal of ZsGreen (TAKARA) and transduced by lentivirus vector LVSIN-IRES-puro, and LgBiT was transduced by lentivirus vector LVSIN-IRES-Hyg [24]. Equal numbers of ZsGreen-HiBiT-expressing cells and LgBiT-expressing cells were plated on a 96-well plate (1603101, ThermoFisher Science). S protein was transduced by lentivirus vector. At 14 hours after transduction, the culture supernatant was replaced to Opti-MEM (ThermoFisher Science), and 25 μL of diluted Nano-Glo live solution (Promega) was added into the each well. After mixing, the relative luminescence was measured by Ensight (Perkin Elmer).

### ELISA assay for direct interaction of recombinant ACE2 and S protein

Diluted recombinant S protein (2μg/mL) (#RP012383LQ, ABclonal) with PBS was incubated to immobilize on the maxisorp plate (464718, Thermo). After blocking with 1% BSA/PBS, 1 μg of K-874A diluted in 1%BSA/PBS was incubated to bind to S protein. Then, serial diluted recombinant ACE2 (from 1 μg/ml to 0.06 μg/mL) (ab151852, abcam), subsequent rabbit anti ACE2 antibody (HPA000288, Atlas Antibodies) and HRP conjugated anti rabbit IgG (7074S, Cell signaling) were incubated. To detect K-874A binding to immobilized S protein, serial diluted K-874A (1 μg/ml to 0.002 μg/ml) in 1%BSA/PBS was incubated, and K-874A which bound to S protein was detected by HRP-conjugated anti VHH antibody (#128-035-232, Jackson immuno Research). Between each incubation, wells were washed with PBST. To color, o-phenylenediamine dihydrochloride (p6662, Merck) was used as a substrate of HRP and reaction was stopped 2M H_2_SO_4_. Signaling was quantified by Ensight (Perkin Elmer).

### Immunofluorescence staining of SARS-CoV-2-infected cells

VeroE6/TMPRSS2 cells were infected SARS CoV-2 and fixed at 24 hours after infection. After treatment with 0.05% Triton-X and 3% BSA/PBS for permeabilization and blocking, cells were incubated with anti-NSP8 (5A10, GeneTex) or anti-dsRNA (rJ2, Merck) for an hour and subsequently with Alexa 568-conjugated goat anti-mouse IgG (A110301, Thermo Fisher) and Hoechest for an hour at room temperature. Images were taken by BZX800 (Keyence).

### Infection to Alveolar organoid and VHH treatment

Human alveolar-derived organoids were established[15] (Ebisudani T et al. submitted). Briefly, 3D cultured organoid compound in Matrigel was treated with cell recovery solution to remove the cells from the Matrigel. The cells were infected with SARS-CoV-2 (MOI 5) for 1 hour at 37°C, washed to remove unbound virus, and compounded into Matrigel. After the compete medium solidified, VHH (15 μg/mL) was added or not. Culture supernatants were collected at each time point and preserved at −80°C.

### VHH treatment of SARS CoV-2-infected Syrian hamsters

2×10^3^ TCID_50_ of SARS-CoV-2 was nasally inoculated to 7 weeks age of Syrian hamsters. Then 3 mg of K-874A was nasally administrated at just before, 1 day or 1 day and 2 days after virus inoculation. The weight of each hamster was measured every day up to 4 days after inoculation. Lung tissues were homogenized at 4 days after inoculation, and RNA was extracted with Nucleospin RNA (TAKARA) for qRT-PCR.

### Quantitative real-time PCR for inflammatory cytokines

Primers and probes for hamsters were designed as reported[25]. The information of primers and probes is shown in Table 1. qRT-PCR experiments were performed on Light Cycler 96 (Roche) with Luna Universal One-Step Rt-qPCR & Probe One-Step RT-qPCR Kits (NEB, USA), according to manufacturer’s instructions. Briefly, 50 ng of RNA was used for each reaction as template. The final concentrations of each test primer and probe set for target gene were 0.4 and 0.2 μM, respectively. The final concentrations of the internal control primer and probe set were 0.2 and 0.1 μM, respectively. Cycling conditions were as follows: 10 min at 50°C (initial reverse transcription), 20 sec at 95°C (inactivation and initial denaturation), and 40 cycles of 3 sec at 95°C, followed by 60 sec at 60°C per cycle.

**Table 1.**
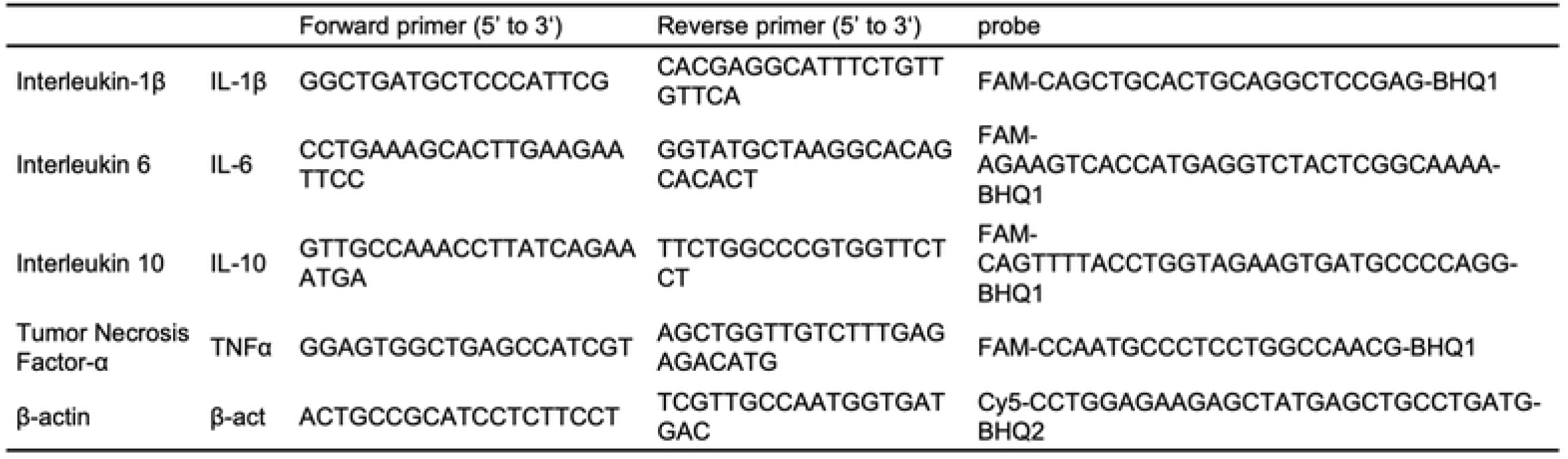
Detailed sequence information of the primers and probes for qRT-PCR

### Cryo-Electron Microscopy

S protein trimer of SARS-CoV-2 (# SPN-C52H9, Acrobiosystems, 600 μg/mL) and K-874A (11 mg/mL) were mixed at 10:1 and kept at room temperature for one day. Aliquot (2.5 μL) of the sample was placed onto a holey-carbon copper grid (Quantifoil Micro Tools, R 1.2/1.3) previously glow-discharged in a plasma ion bombarder (PIB-10, Vacuum Device Inc.). The grid was then blotted (blotting time: 3.5 sec, blotting force: 7) and plunge-frozen at a condition of 95% humidity and 4°C by using a Vitrobot Mark IV (Thermo Fisher Scientific). The cryo-EM data were acquired with a Titan Krios at 300 kV (Thermo Fisher Scientific) and a Gatan K3 camera (Gatan Inc.) at a nominal magnification of 64,000, corresponding to 1.11 Å per pixel on specimen. Each micrograph was recorded as a movie of 58 frames at the total dose of approximately 40 electrons per Å^2^. A GIF-quantum energy filter (Gatan Inc.) was used with a slit width of 20 eV to remove inelastically scattered electrons. Individual movies were subjected to per-frame drift correction by MotionCor2 [26]. The contrast transfer function parameters of each micrograph were estimated using CTFFIND4 [27] and the flowing 2D and 3D classification, 3D refinement, and local resolution calculation were performed with RELION3.1 software [28]. Particles were selected from 6,984 micrographs and the final 3D reconstruction was computed with 115,297 particles. The final map resolution was 3.9 Å (gold standard FSC criterion) by imposing C3 symmetry. Using the “relion_particle_symmetry_expand” command [28], 3 subunits in each particle image were independently compued. The generated 3 × 115,297 (345,891) sub-particle images were used for the 3D reconstruction. A density including K-874A, RBD, and NTD was extracted using UCSF Chimera [29]. The extracted density was used to generate the mask using “Mask creation” in RELION 3.1. The images containing K-874A, RBD, and NTD were subtracted from the symmetry-expanded images using “Particle subtraction” in RELION3.1. The prepared mask was used for subtraction. These sub-particles were subjected to 3D classification without shift and rotation, and the sub-particle images (143,647 particles) of the selected good classes were used for 3D refinement by imposing C1 symmetry. The resolution of the subtracted map was estimated to be 5.0 Å based on a gold standard FSC. The entire procedure is summarized in Supplementary Fig 1 and Table 1.

The SWISS-MODEL [30] server was used to generate homology models of K-874A, and S protein monomer of SARS-CoV-2 (SPN-C52H9) using the atomic model of VHH-72 (PDB ID: 6WAQ) and 2019-nCoV S protein (PDB ID: 6VSB) as a template, respectively. RBD (Pro330-Pro521) and NTD (Ala27-Ser305) were extracted from the homology model, respectively. The homology models of K-874A, RBD and NTD were rigid-body-fitted into the map using the COOT [31] and then refined using PHENIX [32].

### Ethics

Animal experiments were approved by the President of Kitasato University through the Institutional Animal Care and Use Committee of Kitasato University (20-011), and performed in accordance with the Guidelines for Animal Experiments of Kitasato University.

---

**Supplementary table 1.**

| Data Collection |  |
| --- | --- |
| Electron microscopy | Titan Krios |
| Camera | Gatan K3 |
| Voltage | 300 kV |
| Magnification | 64,000 |
| Calculated pixel size | 1.11 Å |
| Exposure time (s) | 5.6 |
| Dose per second (e-/Å<sup>2</sup>/s) | 7.14 |
| Electron dose | 40 electron/Å<sup>2</sup> |
| Number of frames | 58 |
| Defocus range | 0.5 - 2.5 µm |
| Image Processing: Whole structure of S protein trimer and K-874A |  |
| Frame alignment | MotionCor2 |
| CTF estimation software | CTFFIND 4.1.5 |
| Number of micrographs | 6,984 |
| 3D map reconstruction software | Relion 3.1 |
| Initial number of particles | 800,000 |
| Particles contributing to final map | 115,297 |
| Applied symmetry | C3 |
| Applied B-factor | -146 Å<sup>2</sup> |
| Global resolution (FSC = 0.143) | 3.9 Å |
| EMDB number | EMD-XXXXX |
| Image Processing: Focused 3D refinement on K-874A, RBD and NTD |  |
| 3D map reconstruction software | Relion 3.1 |
| Initial number of particles | 345,891 |
| Particles contributing to final map | 143,647 |
| Applied symmetry | C1 |
| Applied B-factor | -470 Å<sup>2</sup> |
| Global resolution (FSC = 0.143) | 5.0 Å |
| EMDB number | EMD-XXXXX |
| Model Building of K-874A, RBD and NTD |  |
| Homology model generation software | SWISS-MODEL |
| Rigid body fitting software | COOT |
| Auto-refinement software | Phenix |
| Number of residues built | 116 (K-874A), 191 (RBD), 278 (NTD) |
| R.m.s. deviation (bonds) | 0.006 |
| R.m.s. deviation (angles) | 1.123 |
| Ramachandran favored | 84.17 % |
| Ramachandran allowed | 15.66 % |
| Ramachandran outliers | 0.17 % |
| Rotamer outliers | 0.20 % |
| C-beta outliers | 0 |
| Clash score, all atomsC-beta | 8.88 |
| PDB ID | XXXX |

